# A gene with a thousand alleles: the HYPer-variable effectors of plant-parasitic nematodes

**DOI:** 10.1101/2023.10.16.561705

**Authors:** Unnati Sonawala, Helen Beasley, Peter Thorpe, Kyriakos Varypatakis, Beatrice Senatori, John T. Jones, Lida Derevnina, Sebastian Eves-van den Akker

## Abstract

Pathogens are engaged in a fierce evolutionary arms race with their host. The genes at the forefront of the engagement between kingdoms, including effectors and immune receptors, are part of diverse and highly mutable gene families. Even in this context, we discovered unprecedented variation in the HYPer-variable (HYP) effectors of plant-parasitic nematodes.

We discovered single effector gene loci that can harbour potentially thousands of allelic variants. These alleles vary in the number, but principally in the organisation, of motifs within a central Hyper Variable Domain (HVD). Using targeted long-read sequencing, we dramatically expand the HYP repertoire of two plant-parasitic nematodes, *Globodera pallida* and *G. rostochiensis,* such that we can define distinct sets of species-specific “rules” underlying the apparently flawless genetic rearrangements. Finally, by analysing the HYP complement of 68 individual nematodes we made the unexpected finding that despite the huge number of alleles, most individuals are homozygous. Taken together, these data point to a novel mechanism of programmed genetic variation, termed HVD-editing, where alterations are locus-specific, strictly governed by rules, and can theoretically produce thousands of variants without errors.

## Introduction

Pathogens are engaged in a fierce evolutionary arms race with their host. Often, and on both sides, the genes at the forefront of the interaction are part of large, diverse, and highly mutable gene families. Pathogen effectors, and host resistance genes, can be part of some of the most numerous gene families in their respective genomes (Tang et al. 2022). Effectors are typically more diverse than other genes in pathogen genomes (Eves-van den Akker et al. 2016), and in some cases are encoded by parts of the genome that mutate rapidly (Rouxel et al. 2011).

Like many pathogen effector genes, HYPs encode nematode proteins that are secreted into the plant, are necessary for infection, have no known homologues, and do not encode any recognised PFAM domains (with the exception of an N-terminal signal peptide for secretion) (Eves-van den Akker et al. 2014). HYPs, however, are highly unusual even among effectors. To date, 75 unique genomic sequences across three subfamilies (HYP1*, n=41;* HYP2*, n=12;* and HYP3*, n=22; average length* 888 bp) have been cloned. Irrespective of subfamily, all cloned HYPs share two continuous strings of coding sequence that are ∼95% identical between genes (410 bp at the 5’ and 94 bp at the 3’). Subfamilies are distinguished primarily, although not exclusively, by a subfamily-specific hyper-variable “domain” (HVD) of unknown function, between the conserved regions. Using HYP1 subfamily members as an example, the HVD can encode up to four “motifs”, some of which themselves contain variable single- or di-amino acid residues (1.1, 1.2, 1.3, and 1.4). Ignoring the variability within each motif, the 41 unique HYP1 genes contain 17 different organisations of these motifs, with little observable pattern. HYP3s show similar levels of variation (although their HVD contains a different number and organisation of an entirely different set of “motifs”), while HYP2 lacks the HVD domain altogether.

Regardless of organisation, the entire HVD is transcribed as a single exon, in frame with the flanking conserved regions: the variation described is genomic, rather than being the result of alternative splicing. Most of the variation within this domain is not just differences in the number of repeats of a particular motif, but differences in the organisations of motifs, which themselves can vary in the sequence that encodes them.

Despite that 75 unique HYP genes have been cloned from a population, no individual nematode genome encodes them all. Individuals vary in the types of effector subfamilies that their genomes encode, and individuals vary in their overall number of HYP genes across an order of magnitude. Importantly, this is not several copies of the same gene, but several different HYP genes that vary in the organisation of their respective HVD (Eves-van den Akker et al. 2014). To the best of our knowledge, there is no known mechanism that can account for both the hyper-variable domain organisations and the apparent gene number variation spanning an order of magnitude between sisters of the same population.

The genetics underlying HYP effectors has remained elusive because previous genome sequencing attempts pre-dated the discovery of HYPs, and used a combination of 76 bp and 100 bp Illumina reads that are shorter than either HYP conserved region (5). To understand the genetic basis of variability and stability in the HYP effectors, we sequenced and assembled long-read genomes of two related species. Strikingly, we found single gene loci for HYP1 and HYP3, indicating that the bewildering diversity of HYPs in fact represents one of the largest allelic series of any organism described to date: conservatively close to a thousand alleles. We use the expanded HYP repertoire to define the rules underlying HYP “editing” in sufficient detail to simulate HVDs *in silico*. Finally, by analysing the HYP complement of 68 individual nematodes we found that despite the huge number of alleles, most individuals are homozygous. Hypothetical solutions which explain the juxtaposition of these apparently contradictory phenomena are discussed, perhaps pointing to the underlying cause of the diversity itself.

## Results

### HYP variation is allelic

To investigate the genetic basis of HYP variation, we needed to first determine the genomic neighbourhood of HYP effectors in individual nematodes. Due to the microscopic size of each animal, we employed single molecule read sequencing (Oxford Nanopore and PacBio) of a population. While we cannot be certain any two reads came from the same individual (even if they overlap perfectly), we can be absolutely certain that each read came from a single individual. By assigning long HYP-containing single DNA molecules from individuals onto a highly contiguous consensus assembly, based primarily on the non-HYP surrounding sequence, we were able to determine the HYP content of individual nematodes.

Therefore, to provide the necessary high-quality consensus assemblies, the genomes of *Globodera pallida* and *Globodera rostochiensis* were re-sequenced and assembled (See methods and Table S1). To identify HYP-containing long reads from single *G. pallida* individuals, a Hidden Markov Model (HMM) for each HYP subfamily. These modelsw were trained the 75 unique cloned HYP sequences available in NCBI (GenBank accession numbers KM206198 to KM206272 (Eves-van den Akker et al. 2014). The parameters were iteratively optimised to accurately identify reads from each subfamily while still maintaining sensitivity for discovering reads with new HYP variants. A total of 251, 334 and 339 unique reads were identified for subfamilies 1, 2 and 3 respectively.

Strikingly, no single read contained more than one HYP, with the exception of HYP2-containing reads. All HYP-containing reads were mapped, in their entirety, to the *de novo* assembly revealing just two, otherwise unremarkable, loci (Figure 1): HYP1-containing reads mapped to *G. pallida* Newton scaffold 46, while HYP2 and HYP3-containing reads mapped to *G. pallida* Newton scaffold 8. Upon closer inspection and comparison to related isolates and species, a broadly conserved arrangement and the ancestral state in the *Globodera* was revealed - a single gene locus for HYP1, and a single gene locus for HYP3 approximately 30 kb away from a two or three-gene locus for HYP2 (Figure 1 and Figure S1).

**Figure 1.**
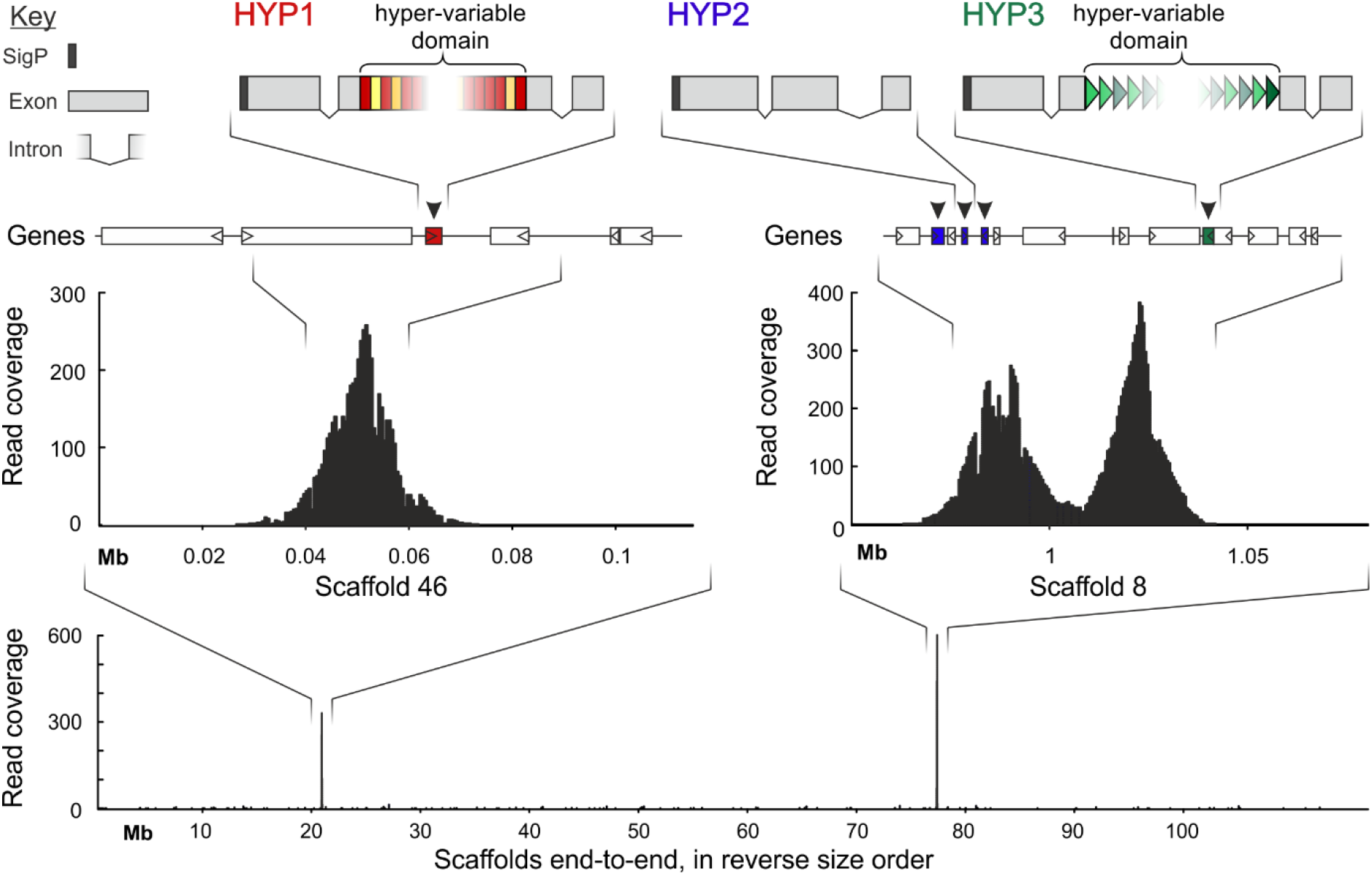
HYP variation is allelic. Schematic of representative HYP1, HYP2, and HYP3, to scale. Hypervariable Domains (HVDs) are indicated by coloured blocks (rectangles in shades of reds for HYP1 and triangles in shades of green for HYP3) within the middle of the middle exon (grey blocks). Location of HYPs and adjacent genes is shown below. Read coverage (black bars) for HYP-containing reads is shown for the HYP1 locus (Scaffold 46, left), and the HYP2 and HYP3 adjacent loci (scaffold 8, right). Substantive coverage is not present on any other scaffolds (bottom).

Taken together, these data show that, strikingly, all of the HYP1 and HYP3 variation observed (which encompasses the dominant majority of all HYP variation) is allelic.

### The rules of HYP variation

To date, four degenerate motifs were discovered for HYP1, and 2 for HYP3, based on Sanger sequencing of cloned HYPs (Eves-van den Akker et al. 2014). To determine a fuller extent of HYP variants, and the frequency in the population of each, a CRISPR enrichment protocol was developed to selectively enrich for the HYP1 and HYP3 loci, providing thousands of unbiased nanopore reads across each locus (Figure 2). To characterise the variants within a *G. pallida* population, a fraction of the available nanopore reads (the top ∼130 highest quality reads (Phred >17)) were used in an iterative approach to, as far as possible, classify the HYP motifs within each read (Figure 2A and S2).

**Figure 2.**
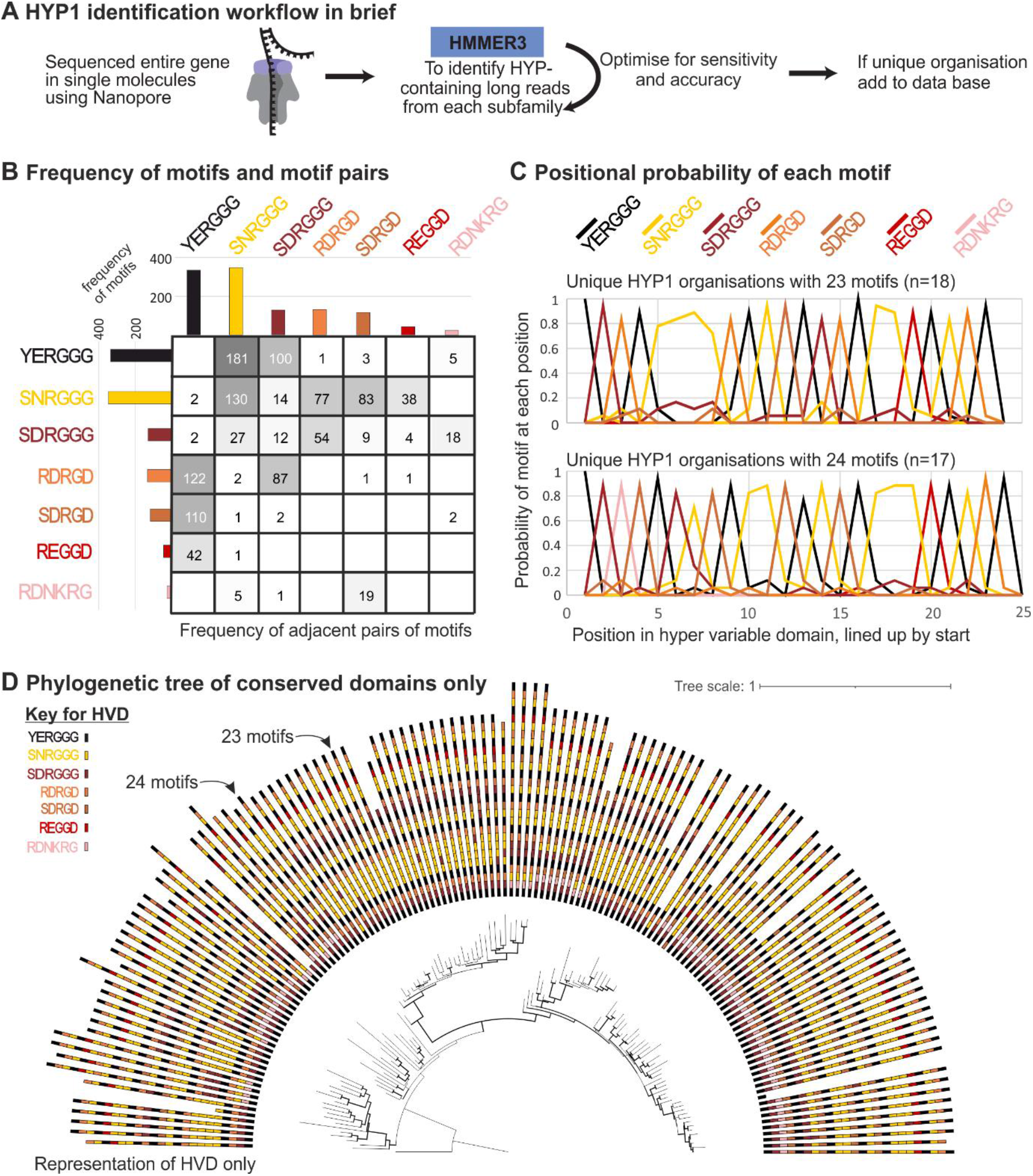
There are rules underlying *G. pallida* HYP1 variation. A) Brief overview of HYP1 identification workflow (see Figure S2 for full). B) Frequency of motif pairs within HYP1 HVDs. Each motif is shown with amino acid sequence, y axis of matrix is position n and x axis is position n+1 (YERGGG is never followed by YERGGG, but it is followed by SNRGGG). C) The positional probability of each motif at each position for HYP1s with HVDs containing 23 (top) and 24 (bottom) motifs. D) A phylogenetic tree inferred from the alignment of the non-HVD domains only, displayed with their corresponding HVDs in the outer semi-circle (colour coded by motif).

In so doing, we were able to re-classify degenerate motifs based on their nucleotide sequence. For example, motif 1.1 ([Y|S][N|E|D]RGGG) can be readily divided into deduced amino acid sequences 1.1.a (YERGGG), 1.1.b (SDRGGG), and 1.1.c (SNRGGG) - Figure S2). The following lists all motifs that can be reliably distinguished in this way in *G. pallida*: 1.1.a (YERGGG), 1.1.b (SDRGGG), 1.1.c (SNRGGG); 1.2.a (RDRGD), 1.2.bc (SDRGD/SDRGE); 1.3 (RDNKRG); 1.4 (REGGD).

We found that the number and organisation of the motifs within the HVD are clearly non-random. Most unique HYP1s contain 23 or 24 motifs within their hyper-variable domain, although this varies greatly (from 1 to 27 motifs). Some motifs occur more frequently than others within unique HYP1s within the entire population (Figure 2B), or within an individual HVD, although this varies greatly: motif 1.1.b is 10 times more common than motif 1.4 overall, and in one case eight times more common within a HVD.

Interestingly, patterns are clearly evident in the organisations of the motifs. Not all possible adjacent motif pairs were observed, and many of those that were observed were not reciprocal: 1.4 (REGGD) was almost always followed by 1.1.a (YERGGG), but 1.1.a was never followed by 1.4. Exceptions include 1.1.b (SDRGGG) and 1.2.a (RDRGD) which were regularly observed following one another. Strikingly, the most common pair (1.1.a (YERGGG) followed by 1.1.c (SNRGGG)) occurs 180 times more than the least common pair (1.2.bc (SDRGD/SDRGE) followed by 1.1.c (SNRGGG)). Homopolymers were almost never observed: only motifs 1.1.c (SNRGGG) and 1.1.b (SDRGGG) occur in homopolymers, with four 1.1.c being the longest observed to date. Of all possible organization of motifs in triples (343) we only observed 75, and the distribution is similarly skewed. The top 20 most common triples include all motifs, and account for 85% of all observed triples, and yet are insufficient to build most HYP1s HVDs with a variable domain >20 (31 out of the 43 HVDs of length longer than 20). This suggests that the skewed distribution of observed triples in the population is not due simply to differences between HYPs, but rather that most individual HVDs are characterised by common higher-order combinations of motifs interspersed with rare higher-order combinations of motifs.

Zooming out further, patterns coalesce. The HYP1 HVD almost always started (i.e. 54 out of 55 times) and ended (i.e. 47 out of 55 times) with motif 1.1.a (YERGGG). More specifically, HYP1 hypervariable domains often end in the following block of five motifs: 1.4 (REGGD), 1.1.a (YERGGG), 1.1.c (SNRGGG), 1.2.a (RDRGD), 1.1.a (YERGGG). The probability of each motif in each position (Figure 2C), together with the analysis of adjacent pairs and triples, revealed clear rules to the organisation of motifs within the HVD at the scale of the HVD itself. Interestingly, a phylogenetic tree inferred from the non-HVD domains of HYP1s (Figure 2D) grouped certain categories of HVDs. This suggests that although the non-HVD is highly conserved it somehow carries information about the structure of the HVD itself.

Based on just three rules: 1) possibility of a given motif to follow another one (Figure 2B); 2) probability of motif at each position (Figure 2C); and 3) known starts and ends - we can simulate HYP1s with >20 motifs that look “normal”. When we simulate a million HYP1 HVDs *in silico*, we re-identify most known organisations and extend the theoretical limit to >73,000 unique variants. This suggests that we have developed a sufficient understanding of the rules of HYP variation to recreate known HYPs *in silico*, and thereby provide a theoretical limit in the tens of thousands, based on extant examples. In contrast, these rules do not apply to the HYP1s of the closely related species, *G. rostochiensis* (GrHYP1s). GrHYP1s are shorter (typically 11 or 13 motifs), composed of an almost entirely different set of motifs, and seem to obey a very different set of rules, but obey rules nonetheless (Figure S3).

### HYP variation is conservatively estimated to exceed a thousand alleles

The theoretical limit of HYP1 variation will massively exceed the true limit because reality is constrained in a multitude of ways that the simulation is not. To estimate the number of alleles in the population, we adopted a population genetics approach. Using the ∼130 highest quality (Phred>q17) nanopore CRISPR-enrichment reads for each species, we identified 55 unique GpHYP1 variants and 90 unique GrHYP1 variants. Not all variants were equally represented within the populations sequenced. In fact, the distribution was extremely skewed, roughly following Zipf’s law: the most common sequence is about twice as common as the next, which itself is about twice as common as the next, etc.. The top two most common HYP1s account for 3.6% of unique variants but 50% of all reads, while at the same time, 47 variants each only occur once and account for 85% of unique variants but only 35% of the reads (Figure 3). A similar pattern is observed for HYP1s of *G. rostochiensis* (albeit with an even greater proportion of unique variants - 93% of unique variants only occur once (84/90), representing 67% of reads).

**Figure 3.**
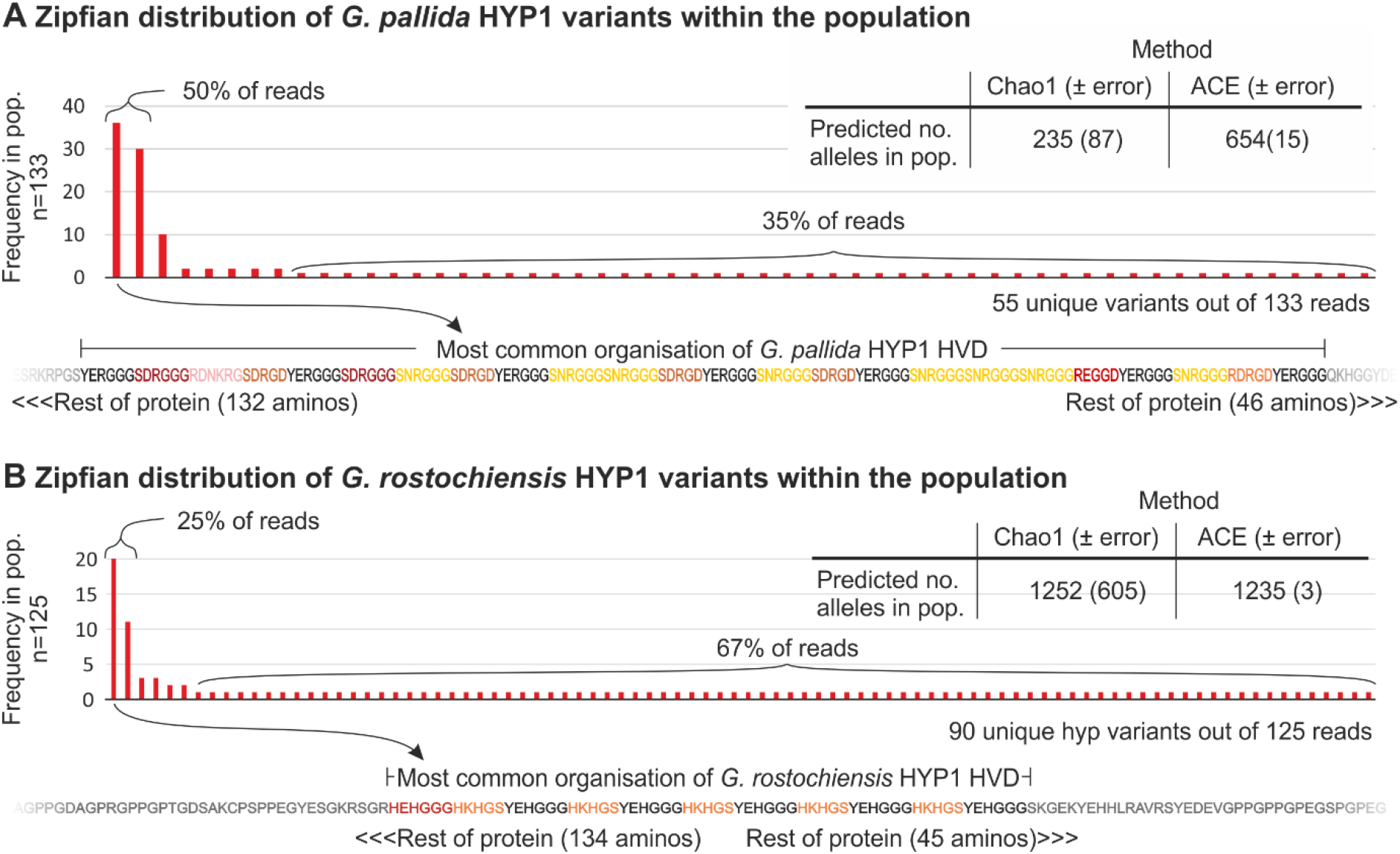
Frequency distribution and population size estimates of HYP1 alleles. A *G. pallida* HYP1 and B *G. rostochiensis* HYP1 alleles. The graphs show the observed occurrences of each unique HYP1 variant when sampling a population (n= 133 and 125 for *G. pallida* and *G. rostochiensis* respectively). Inset are two independent species abundance estimates for the total number of alleles in the population, based on the sampling. Below shows the HVD of the most common variant for each species.

We can extrapolate to the total number of alleles in the population by using the frequency distribution of unique sampled alleles in classical species richness calculations. Using the 133 reads alone, the total number of *G. pallida* HYP1 alleles is estimated to be 235 and 654 using Chao (O’hara 2005) species estimation or the Abundance-based Coverage Estimator (ACE, (O’hara 2005; Chiu et al. 2014)), respectively (Figure 3A). The total number of predicted alleles for HYP1 in *G. rostochiensis* is even higher; 1252 and 1235 using Chao species estimation or the ACE, respectively (Figure 3B). HYP3 alleles show a similar distribution but are generally less variable than HYP1s in *G. pallida*: 18 unique out of 130, estimated population size of 31 and 48 for Chao and ACE respectively (Figure S4). Given that the bulk of HYP variation is HYP1 variation, HYP1s are the focus of this manuscript.

Given that we: 1) analysed a single population of a global pathogen; 2) knowingly ignored all variation outside the HYP domain (Figure 2D); and 3) were conservative in predicting variants in general - these estimates are likely underestimates. Taken together, we predict that global HYP1 variation likely exceeds a thousand alleles per locus.

### Apparent non-mendelian inheritance of HYP alleles

To ascertain the contribution of each individual nematode to the overwhelming diversity of HYPs at the population level, we adapted a PCR protocol to reliably function for two amplifications from a second stage juvenile nematode (J2). From each of 68 individual J2s the HYP1 and HYP3 loci were amplified (with unique barcode pairs appended to the primers), pooled and sequenced using PacBio HiFi. In parallel a series of control amplifications were carried out from known homozygous or heterozygous pools of plasmids containing HYPs.

Consistent with the nanopore data, we found that few HYP variants dominate and that many variants occurred very rarely. As is common for a metagenetic approach in general (Eves-van den Akker et al. 2015), within each barcode pair, even homozygous controls, several different HYP alleles could be identified. These could be explained by multiple HYPs in the starting DNA, cross contamination, miss characterisation of barcodes, and/or sequencing errors (in the gene or the barcode). The controls allowed us to establish a threshold to confidently distinguish between homozygous and heterozygous starting material. As expected, known homozygous controls have most (>90%) of their reads corresponding to a single HYP variant. Heterozygous controls have about half (approx. 40%) of their reads corresponding to a single HYP variant. Analysing the proportion of reads for each barcode pair contributed by the most common allele allowed us to determine whether the underlying control sample was homozygous or heterozygous, without error (Figure 4A). We could not, however, reliably distinguish unequal ratio controls (1:100, 1:500, 1:1000, 1:2000, and 1:5000) from homozygous controls (Figure S5). Therefore, for reads from individual animals of unknown zygosity, we established a threshold of >80% for a likely homozygous sample.

**Figure 4.**
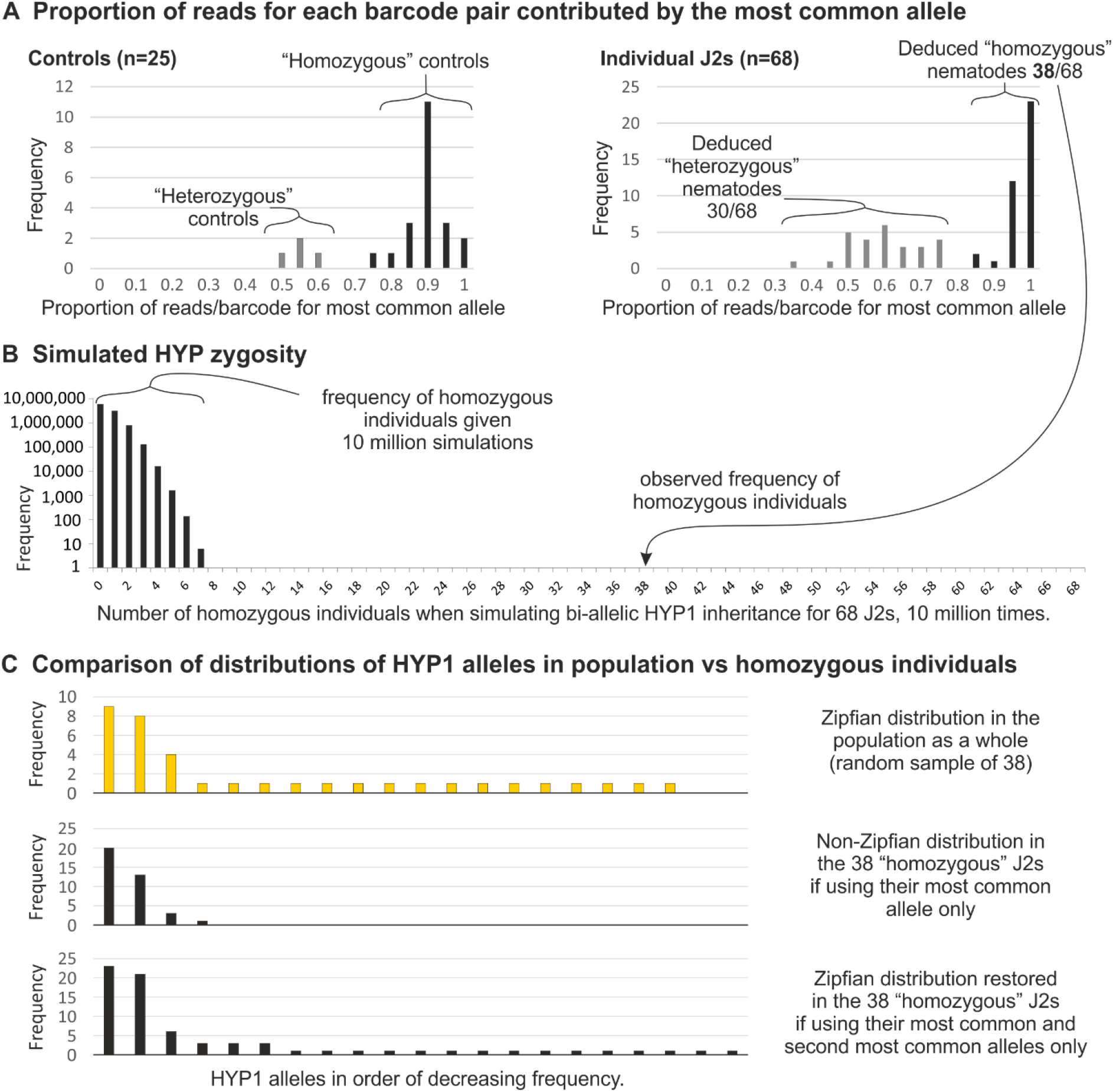
Apparent non-mendelian inheritance of HYP alleles. A) Frequency distributions showing the proportion of reads for each barcode pair that is contributed by the most common allele, for the 25 control samples (left) and the 68 J2 samples (right). Grey bars indicate “heterozygous” samples (known for deduced) and black indicate “homozygous” samples (known or deduced). B) Frequency of homozygous individuals when simulating bi-allelic HYP1 inheritance for 68 J2s, 10 million times. C) Frequency distribution of alleles in: the population (top, yellow); the 38 “homozygous” J2s if using their most common allele only (middle, black); and the 38 “homozygous” J2s if using their most common and second most common alleles only (bottom, black).

Computing the zygosity of individual J2s revealed a surprising anomaly: most individuals were homozygous (Figure 4A). This result was surprising because the probability of being homozygous decreases as a function of the number of alleles in the population, which we estimated to be over a thousand. Known heterozygous controls do not show homozygosity using these criteria, so we reasoned this was not an artefact of PCR “selecting” one allele over the other. To calculate the improbability of this result, we made the conservative assumption that the known 55 HYP1 alleles are all that exist, and empirically derived the probability of 38/68 individuals being homozygous to be substantially less than one in 10 million: after 10 million simulations of selecting two parental alleles from the population for each of 68 nematodes, the highest number of homozygous individuals was 8/68, which occurred once (Figure 4B). HYP3 had a similarly large proportion of homozygous individuals (19/45), but seems to be independent from the HYP1 locus (i.e. individuals homozygous for HYP3 are not also preferentially homozygous for HYP1, Figure S6).

Importantly, dominant alleles in those homozygous individuals are not diverse enough to explain the distribution of alleles in the population (Figure 4C). Analysing the most common allele in each of the 38 HYP1 homozygous individuals revealed just four variants in total, which include the top three most common alleles from the nanopore sequencing (36/133, 30/133, and 10/133) and only one rare allele (1/133). Combining the second most common alleles in all 38 “homozygous” individuals reveals 17 additional variants and 15 of which only occur once in this set. Including 1st and 2nd alleles in “homozygous” individuals is sufficient to recreate the Zipfian distribution observed in the population data.

## Discussion

Although we have been extremely conservative at each step when constructing estimates of HYP diversity, an effector with one thousand alleles is unprecedented in pathology, and perhaps in genes in general. HYP allelic variation even exceeds the most widely recognised hypervariable genes, immunoglobulins. While thousands of immunoglobulin variants have been characterised, and theoretically millions of variants are possible (1×10^7 in humans) (Schroeder, 2006), V(D)J diversity is achieved by recombination of segments that exist as multiple copy arrays on the chromosome. By contrast, HYPs are single copy loci with potentially thousands of alleles.

We can confidently rule out sequencing errors contributing in whole, or even in small part, to both the diversity and the nature of the diversity because: we have used three sequencing technologies (Nanopore, PacBio, and Sanger) each of which reveals HYP variation; two species, each of which show considerable, but different, HYP variation; and importantly we have found two sets of quite different rules, the nature of the variation in *G. rostochiensis* (a pseudo alternating pattern, Figure S3) is largely different to the nature of the variation in *G. pallida* (shuffling of blocks).

Phylogenetics points to an accumulation of HVD alleles, rather than a contraction of multi-copy arrays into a single copy locus, over evolutionary time. The synteny between *G. pallida* and *G. rostochiensis* suggests their last common ancestor had all three HYP families. Whether it had HVDs of today’s diversity is unknown, but given that the HVDs of *G. pallida* and *G. rostochiensis* today are markedly different from one another and appear to obey very different sets of rules (Figure S3), we know that the present day complement of HYP alleles in at least one, but perhaps both, of *G. pallida* or *G. rostochiensis* must have arisen since their divergence (i.e. the last 30 million years). Supporting this hypothesis, the close outgroup *Heterodera schachtii* has two HYP-like genes (Hsc_gene_1517 and Hsc_gene_17937) with yet again entirely different HVD structures that neither resemble HYP1s or HYP3s from either Globodera species (Siddique et al. 2022). Taken together, this suggests a proto-HYP predated the divergence of Heterodera spp. and Globodera spp., which is estimated to be some 100 million years ago.

To understand how HYP diversity evolved, i.e, the pathway from one to one thousand alleles, we need to understand how HYP diversity is created. We intuitively rule out random mutations and subsequent selection because they seem incongruent with the diversity and nature of HYP variants (Figure 2B and C). Similarly, we must rule out mechanisms akin, even in small part, to V(D)J recombination because of: a) genetic capital; and b) imprecision. In terms of genetic capital, V(D)J diversity requires a reservoir of multiple copy arrays on the chromosome that HYPs (Figure 1) do not require. Most likely, the genetic capital used to generate new HYP variants comes from within the HVD itself. In terms of imprecision, V(D)J domains are joined by non-homologous end joining, resulting in imprecise joints that contain added nucleotides, contributing to the diversity (Chi et al. 2020). The source of HYP diversity is apparently flawless. This is evident from extant HYP genes differing in only the middle of the middle exon without disrupting the open reading frame, despite rearrangements of motifs that differ in length. A mechanism of genome editing that is aware of frame, is both unexpected and more akin to guided shuffling/rearranging of the HVD (without inversion), rather than error-prone assembly from a diverse repertoire. We term this phenomenon HVD-editing.

Our primary hypothesis on HVD-editing comes from the striking observations that: i) sequencing a population reveals many rare alleles, and ii) despite potentially thousands of alleles, most individuals appear homozygous for common alleles. We therefore conclude that there must be some mechanism to bias apparent inheritance. Four options are considered: 1) selective mating; 2) abortion of heterozygote zygotes; 3) gene conversion; and 4) post embryonic hypermutation in the soma (Figure 5). Selective mating and zygote abortion seem highly improbable because the number of alleles could easily be greater than the number of nematodes that may infect a single plant. Sex resulting in progeny, in either scenario, is deemed to be sufficiently rare to doom the species to extinction.

**Figure 5.**
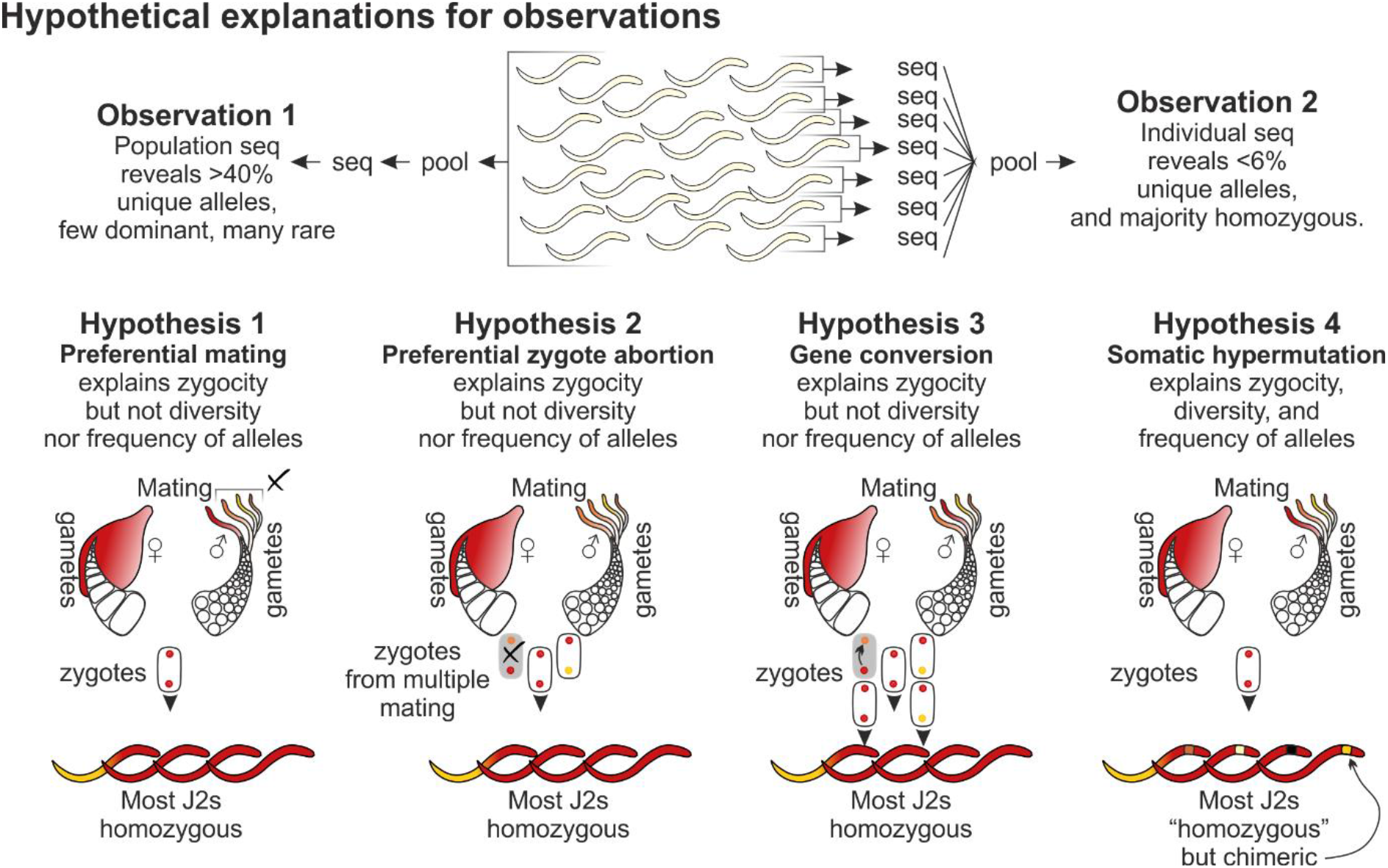
Figure of Hypotheses. The primary observations that: 1) when sequencing a pool of individuals from a population we observe many unique alleles (with a long tail of rare alleles); but 2) when we sequence individuals from that same population and pool their dominant allele per individual we get very few unique alleles (with no long tail of rare alleles). Four potential hypotheses are discussed (Preferential mating, Preferential zygote abortion, Gene conversion, and Somatic Hypermutation). Hypothesis 4, somatic hypermutation, is the most parsimonious explanation as it alone explains all phenomena observed.

The gene conversion hypothesis only explains the improbably high proportion of homozygous individuals, but fails to account for the lack of diversity between said individuals. Consequently, somatic hypermutation remains the only hypothesis we cannot rule out, as it explains all observed phenomena. Specifically, the Zipfian distribution of allele frequency of the population can only be recapitulated with individual whole nematode sequencing if we include the second most common allele in “homozygous” individuals. This may initially seem counterintuitive, but is analogous to what would happen if individual whole humans were sequenced and their immunoglobulin diversity analysed: some rare and re-arranged immunoglobulins, with some “dominant” alleles that have apparently biased inheritance. This observation would only mase sense with the understanding that V(D)J recombination takes place in a small subset of somatic tissues, namely T- and B-cells. If a similar idea, albeit a different mechanism, gives rise to polyzygotic individuals at the HYP locus from a homozygous progenitor, it is tempting to think this would take place in the two cells in which HYPs are expressed, the amphid sheath cells (Eves-van den Akker et al. 2014). This would also elegantly explain the observed ratios of HYP variation within the individual and within the population (Figure 5). A programmed difference between the genomes of the germline and the soma, as seen for immunoglobulins, is also important because it means that the mechanism(s) is(are) still happening today - we would be observing an ongoing process, not cataloguing what evolution has selected from a greater pool of diversity.

If novel alleles within the individual resemble derivatives of the dominant allele, then this would also favour somatic hypermutation over conversion. I.e. the second most common allele in “homozygous” individuals would be related to the most common allele, in ways that the two most common alleles in “heterozygous” individuals would not be related to one another. Indeed, the second most common alleles in “homozygous” individuals are typically shorter than the dominant allele (22/38 J2), whereas for “heterozygous” individuals they are not (2/24). Similarly, not all HYPs have all motifs within their HVD. So we would expect 2nd alleles in “homozygous” individuals, if they were derivatives of the dominant allele, to contain only those motifs present in the dominant allele, but we would not expect the same constraint on the two most common alleles in “heterozygous” individuals. Indeed, 37/38 2nd alleles in “homozygous” individuals contain exclusively motifs present in the dominant allele, whereas the opposite is observed for “heterozygous” individuals (1/24). Taken together, classical Mendelian inheritance of the locus, coupled with somatic hypermutation in the soma, would explain all observed phenomena.

While determining the teleology of phenomena is often challenging, it is particularly so for HYP variation. Programmed variation is relatively rare, but widely distributed across the tree of life including invertebrates and vertebrates (Wang et al. 2017). When and why organisms distinguish between germline and somatic genomes varies, but can include the “silencing” of germline-active genes in the soma (e.g. programmed DNA elimination (Wang et al. 2017; Gonzalez de la Rosa et al. 2021)) or restricting the generation of variation to the soma, to balance adaptability in recognising invading pathogens with genome stability (Chi et al. 2020). HYPs are involved in inter-kingdom interactions, expressed during parasitism, and required for full pathogenicity (Eves-van den Akker et al. 2014), but in this case they are deployed by the invading organism. What aspect of plant-biology requires such extreme diversity in the pathogen is not clear, nor whether it is diversity to overcome some aspect of one host, or the differences between hosts/host species. The implication would be that the allele you are born with, and presumably the one you pass on to your offspring, is not the one you interact with the plant with. Plant-parasitic nematodes face a very uncertain environment, the host might change every year, and one can imagine strategies evolving that maximise “getting through the year”.

Given that pathogens are engaged in a fierce co-evolutionary arms race with their host, it should not be surprising to uncover yet more unusual, but potentially useful, biology (c.f. CRISPR (Jinek et al. 2012); TALENs (Boch et al. 2009); and transgenesis (Joos et al. 1983)). Taken together, our working hypothesis is that there is remarkable and potentially useful biology underlying HVD variation, it is active today, it creates variation from genetic capital within the HVD itself, it does so in a subset of the soma (most likely the cells in which they are expressed), it is precisely guided, and it can theoretically produce thousands of such variants without scars. Future work will test these hypotheses.

## Methods

### DNA extractions

For DNA extractions from a population, a pellet of freshly hatched J2s was flash-frozen in liquid nitrogen. Between 30,000 and 50,000 J2s were used for high-molecular weight (HMW) DNA extraction, resulting in ∼3.5-5µg of DNA. This was scaled up as required to make larger/multiple libraries. Briefly, a pellet of frozen J2s was homogenised in 20 µL of lysis buffer (0.1M Tris at pH 8.0, 0.5M NaCl, 50mM EDTA and 1%SDS) using a micropestle. Additional 140 uL of lysis buffer and 40uL of proteinase K (20mg/ml, Promega, Cat. No. MC5005) were added, and incubated at 55 °C for 18 hours. 10µL of RNAseA (10mg/ml, Thermo Scientific, Cat. No. EN0531) was added, mixed gently and incubated for 5 min at room temperature. Equal volume of phenol/chloroform/isoamyl alcohol was added to the lysed J2s and mixed by rotating on a hula mixer for 15 min. Post centrifugation, the aqueous solution was collected in a fresh tube and the above step was repeated by adding a 10mM Tris-HCl (pH 8.5) buffer to the organic phase. The DNA was further purified by using equal volume of chloroform/isoamyl alcohol (24:1) for one round of back extraction. DNA was precipitated by adding NH_4_OAc (0.75M), glycogen (20 µg) and 2.5 volume of 100% ethanol. The DNA was centrifuged at 4 °C for 20 min. The resulting pellet was washed two times with 80% ethanol, air dried, and resuspended in 10 mM Tris-HC (pH 8.5). DNA in aqueous solution was handled using wide-bore tips and extraction was performed using low-retention microcentrifuge tubes. DNA amount was measured using Qubit fluorometer (Invitrogen) and its purity was assessed by measuring the absorbance ratios using NanoDrop 1000 spectrophotometer (ThermoFisher Scientific).

For DNA from single nematodes, extractions were performed in worm lysis buffer (WLB) (50mM KCl, 10mM Tris (pH 8.3), 2.5mM MgCl_2_, 0.45% NP-40 (IGEPAL), and 0.45% Tween-20) with Proteinase K (20mg/ml, Promega, Cat. No. MC5005) added just before use. Single nematodes were placed in PCR strip tubes containing 5µL of WLB. They were subjected to three rounds of freeze-thaw using flash-freezing in liquid nitrogen. An additional 5µL of WLB containing Proteinase K (12µL of Proteinase K in 88µL of WLB) was added to each tube. The nematodes were then digested by incubating the PCR strips at 65°C for 90 minutes followed by inactivation of Proteinase K at 95°C for 15 minutes in a Thermal Cycler (Applied Biosystems ProFlex PCR System).

### Oxford Nanopore and PacBio Sequencing

Nanopore sequencing libraries were prepared using the Ligation Sequencing kit (SQK-LSK109, Oxford Nanopore) and the associated manufacturer’s protocol (version: GDE_9063_v109_revQ_14Aug2019) with the following modifications: Incubation with and elution from AMPure XP beads (Beckman Coulter, Cat. No. A63882) was extended to 30 minutes each. Incubation for ligating adapters was extended to two hours at room temperature.

Potato cyst nematode DNA appears to block the pores in the flowcells within 12 hours of running on a MinIon, resulting in fewer reads. Hence, when possible, two libraries were prepared and Flow Cell Wash Kit (WSH003/ WSH004, Oxford Nanopore) was used to regenerate the pores after the first run before loading the second library. The second library was stored at 4°C until loading. One flowcell run was used to generate sequencing data for *G. pallida* ‘Lindley’ and three flowcell runs were used for *G. rostochiensis* ‘Ro1’; two of these runs were from DNA size-selected using 15 kbp and 30 kbp cutoff points, respectively. Size selection was performed using BluePippin and the associated PAC20KB kit (Sage Science) according to the manufacturer’s protocol.

For PacBio sequencing of *G. pallida* ‘Newton’, 20Kb shear was performed and size selection was performed using a Blue Pippin (Sage Science) with an 8Kb cut-off point. The library was sequenced on Pacific Biosciences Sequel instrument.

### PacBio Hi-Fi sequencing for barcoded PCR from individual nematodes

Randomised barcodes of 9bp and 10bp length and a minimum Hamming distance of five were generated using the generate-barcodes script from https://github.com/audy/barcode-generator (Supplementary File. 1). PCR using barcoded primers was performed using KOD Xtreme Hot Start DNA Polymerase (Cat. No. 71975-3) using the following reaction mixture: 25µL 2X Xtreme buffer, 10µL dNTPs (2mM), 1.5µL each of forward and reverse primers at 10µM, 6µL of nuclease-free water, and 5µL of single nematode DNA (extracted as described earlier). The PCR was performed using the following conditions: initial denaturation at 94°C for 2 minutes; 40 cycles of 98°C for 10 seconds, 55°C for 30 seconds and 68°C for 1 minute 30 seconds for HYP1 gene (and 2 minutes for HYP3 gene); final extension at 68°C for 7 minutes.

10µL of the PCR product was used to visualise amplification via gel electrophoresis on a 1% Agarose gel containing SYBR™ Green I Nucleic Acid Gel Stain (Thermo Fisher Scientific). The rest of the PCR product was cleaned using Monarch PCR & DNA cleanup kit (Cat. No. T1030L). The cleaned-up PCR products were then pooled by roughly normalising for amounts based on the intensity of their band visualised during gel electrophoresis. ∼4µg of the pooled amplicons were used for PacBio HiFi sequencing on a Sequel II instrument.

### Genome Assembly and Annotations

Genomes from Nanopore reads (*G. rostochiensis* ‘Ro1’ and G.pallida ‘Lindley’) were assembled using wtdbg2 (Ruan and Li 2020). In addition to the preset -x ont, the following additional parameters were used for *G. pallida* ‘Lindley’: -S 1 and -A and the following for *G. rostochiensis* ‘Ro1’: -S 1 -A -L 10000. PacBio subreads generated from four flow cells for *G. pallida* ‘Newton’ were concatenated and error-corrected using Canu (v1.7 (Koren et al. 2017)) with the following additional parameters: correctedErrorRate=0.15 corOutCoverage=300. Canu error-corrected reads were used to generate the final assembly using wtdbg2 with the following parameters: -L 4000 and -p 19.

For all genome assemblies, purge haplotigs pipeline was used to remove duplicated contigs from the primary assembly (Roach et al. 2018). The primary contig assemblies were subsequently assessed for contamination using BlobTools version 1.0 (Laetsch and Blaxter 2017). Briefly, reads were mapped to the assembly using minialign version 0.5.2 (https://github.com/ocxtal/minialign) to determine the coverage of the assembled contigs. The contigs were BLASTn searched against GenBank nt database with taxonomic information in the tabular output. Contigs were then taxonomically classified based on the weight of the BLAST hits. Identified contaminant contigs were removed thus, yielding a contamination free final unpolished contig level assembly. The assembly was further improved using FinisherSC (Lam et al. 2015). SSPACE-LongRead (Boetzer and Pirovano 2014; Lam et al. 2015) was used to scaffold the contigs using the following parameters: -k 1 -o 1000 -l 30 for Nanopore assemblies and -g 500 -l 10 -o 100 -k 1 for the PacBio assembly. Gaps in the assembly were filled using gapFinisher (Kammonen et al. 2019). The scaffolds were further polished and corrected using five rounds of Pilon (Walker et al. 2014) using both raw reads from Nanopore/PacBio sequencing and short paired-end reads from Illumina HiSeq 2000. Following short reads accessions were downloaded from ENA and used for assembly correction (https://www.ebi.ac.uk/): ERR114517 and ERR114518 for *G. pallida* ‘Lindley’ and ERR114519, ERR123958, and ERR114520 for *G. rostochiensis* ‘Ro1’. minimap2 (Li 2018) was used to map nanopore reads and BWA-MEM (Li and Durbin 2009) was used to map the short reads to the genome assembly. https://github.com/peterthorpe5/public_scripts/tree/master/gene_model_testing

Transposon prediction, and hard and soft repetitive genome masking, was performed as described in (Thorpe et al. 2018). Briefly, Repeatmodeler (version DEV) was used to identify repetitive regions. The resulting repetitive elements were masked using RepeatMasker along with RepBaseRepeatMaskerEdition-20170127 models. To additionally identify transposons, TransposonPSI version 08222010 (Haas 2007) and LTRharvest version 1.5.9 (Ellinghaus et al. 2008) from Genometools (Gremme et al. 2013) was used. Finally, bedtools (Quinlan 2014) version 2.27.1 was used to softmask the genome for gene prediction. RNAseq data from the relevant species (Cotton et al. 2014; Eves-van den Akker et al. 2016) was quality (Q30) trimmed using trimmomatic, allowing a minimum read length of 67, and mapped to the final genomes using STAR version 020201 (Dobin et al. 2013) with the following parameters -- outFilterMismatchNmax 7 --outFilterMultimapNmax 5. The resulting bam files were merged, sorted and indexed using samtools. The sorted bam was used to perform a *de novo* genome-guided RNAseq assembly using Trinity version 2.8.4 (Haas et al. 2013) with the additional parameters (--genome_guided_max_intron 15000 --genome_guided_min_coverage 5). The softmasked genome along with the RNAseq mapped .bam file was subjected to unsupervised gene prediction using BRAKER version 2 (Hoff *et al.,* 2015) with the additional Augustus parameters --protein=on --start=on --stop=on --cds=on --introns=on --noInFrameStop=true -- genemodel=complete and the additional BRAKER parameters --filterOutShort --UTR=on. The resulting BRAKER predicted GFF file and the GeneMark-ET (Lukashin *et al.,* 1998) GFF files were passed to Funannotate DOI: 10.5281/zenodo.2604804 with the weighting of 5 and 1 respectively (out of 10). DIAMOND_BLASTP search against Swiss-prot was also given as evidence during the prediction phase. The genome-guided assembly was given to Funannotate which runs PASA, the resulting PASA models were given a score of 6 (out of 10) in the Evidence modeller stage. The update stage of Funannotate refines the introns, exon, start, stops using a combination of PASA and Stringtie (Pertea et al. 2015). Gene calls were tested with https://github.com/peterthorpe5/public_scripts/tree/master/gene_model_testing

### Finding and filtering HYP containing reads to identify HYP loci

Previously cloned HYP genes were used to build an HMM model using the hmmbuild function from HMMR3 for each of HYP subfamily (HYP1, HYP2 and HYP3) (See Supplementary Files. 2-4 for alignments of HYP genes used to build HMM models for each subfamily). HYP containing long pacbio raw reads were identified by using nhmmer3 function in HMMR3. Multiple evalues were tested to ensure that the reads did not overlap between subfamilies. Evalue of 1e-100 was found to be optimal. These reads were then mapped back to the genome using minimap2 (minimap2 -ax map-pb --secondary=no). The aligned reads were converted to a bedgraph coverage track using bamCoverage function from the deepTools package (using -bs 10000). The plots were plotted using the plotBedgraph() function from the Sushi package in R (Phanstiel et al. 2014). Schematic gene maps were drawn using the ggplot2 extension gggenes and then edited in vector graphics editors.

### CRISPR enrichment of HYP1 and HYP3 loci gRNA design and duplex assembly

crRNAs (IDT) were used with tracrRNA (IDT) to form a functional gRNA duplex. FlashFry (McKenna and Shendure 2018) was used to identify and design crRNAs. Guides were designed to flank HYP1 and HYP3 genes for *G. pallida* and *G. rostochiensis* using the PacBio *G.pallida* (Newton) assembly and Nanopore - corrected *G. rostochiensis* (Ro1) assembly. Candidate guides were filtered using these following parameters: dangerous_in_genome = “IN_GENOME=1”, dangerious_GC and dangerous_polyT = NONE and baseDiffToClosestHit >= 3. Candidate guides were then validated using *in vitro* Cas9 cleavage assay, and guides with the highest cleavage efficiency were used for enrichment (Table 1).

**Table 1.**
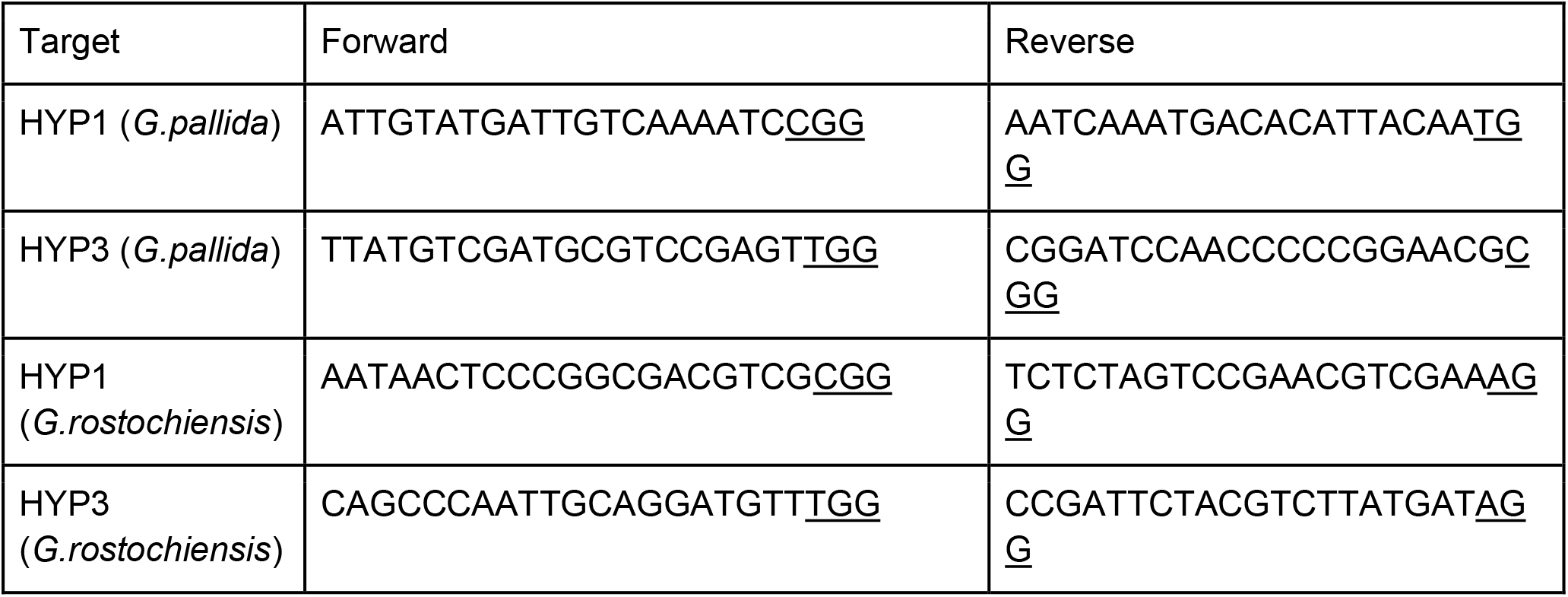
Table of crRNAs used for targeting HYP1 and HYP3.

SQK-LSK109 ligation sequencing kit was used for the library preparation of the test run and SQK-LSK112 ligation sequencing kit was used for the following runs. The test was run on a Flongle. The rest of the runs were run on MinIon flow cells (version R10.4.1). Libraries were prepared by adapting the Cas9-targeted sequencing protocol from Nanopore (version ENR_9084_v109_rev_04Dec2018). Briefly, 0.75µL of each of the cRNAs (100µM in TE, pH 7.5, IDT) (Supplementary Table 1) were pooled together. 2µl of this mixture was added to 2µL of tracrRNA (100µM, IDT) and 6µL of Duplex Buffer (IDT). This was mixed and heated at 95°C for 5 minutes in a Thermal Cycler (Applied Biosystems ProFlex PCR System). 10µL of the annealed cRNA-tracrRNA pool was added to 10µL of NEB CutSmart buffer, 79.2µL of nuclease-free water and 0.8µL of Cas9 nuclease (62µM, IDT Cat. No. 1081059). The ribonucleoprotein complexes (RNPs) were allowed to form by incubating the tube at room temperature for 30 minutes. The genomic DNA (extracted as described above) was dephosphorylated by adding 1.5µL of rSAP (NEB, Cat. No. M0371S) to 25.5µL of HMW DNA and 3µL of NEB CutSmart Buffer. The tube was mixed and incubated at 37°C for 1 hour followed by inactivation of rSAP at 65°C for 10 minutes. 10µL of the Cas9 RNPs were added to the dephosphorylated DNA along with 1µL of freshly prepared 10mM dATP (NEB, Cat. No. N0440S) and 1µL of Taq polymerase (NEB, Cat. No. M0273). This was incubated at 37°C for 1 hour followed by 72°C for 5 minutes in a Thermal Cycler to cleave the DNA by Cas9 and dA-tail the target DNA. This was followed by adapter ligation, clean-up and flowcell loading following the protocol for SQK-LSK112 sequencing kit (Version GDE_9141_v112_revC_01Dec2021) with the following modifications: Incubation with and elution from AMPure XP beads (Beckman Coulter, Cat. No. A63882) was extended to 30 minutes each. Incubation for ligating adapters was extended to two hours at room temperature. Sequencing reads were basecalled using Guppy v4.4.1.

### Enrichment plots

Coverage for HYP1 and HYP3 loci was plotted using GenomicRanges and GenomicAlignment in R, adapting the script from Giltpatrick et al (Gilpatrick et al. 2020).

### Motif analysis for HYP1 and HYP3 genes from Nanopore and PacBio HiFi sequencing

A custom pattern-matching script was developed to retrieve HVD sequences from Cas9-enriched and PacBio HiFi using the Biostrings and stringr packages in R. All the scripts and notes on how to use them can be found at https://github.com/unnatisonawala/HYPervariable_HYPs In case of Cas9-enriched Nanopore sequencing from a population of J2s, the reads were filtered additionally filtered for Q>17 for *G. pallida* and Q>15 for *G. rostochiensis* using NanoFilt (De Coster et al. 2018) and manually curated for motifs not identified by the pattern-matching script due to Nanopore errors.

Species richness for HVD variants were calculated using estimateR function from the vegan (Dixon 2003) package in R.

### Zygosity

To empirically derive the probability of 38/68 individuals being homozygous in spite of the large number of apparently available alleles in the population, a custom python script was written (https://github.com/sebastianevda/HYP/tree/main/Zygosity). The script, based on a population of alleles, randomly selects 2 "parents" for each of 68 individuals, and computes zygosity. The population of alleles used was HVD domains of all 133 HYP1 nanopore reads (i.e. maintaining the frequency of each unique allele). Ten million iterations were performed.

### Simulating HYP HVDs *in silico*

All nanopore-derived unique HYP HVDs (55) were lined up by their start position, and the probability of each motif occurring at each position was computed (including no motif, i.e. the end of an HVD). A custom python script (https://github.com/sebastianevda/HYP/tree/main/Simulate_HVDs/Compute_HVDs_based_o n_positional_probability_and_known_pairs.py) was written to generate HVDs motif by motif, based on the probability of a given motif at that position (probability of end of HVD at position 29 is 1), provided that the selected motif is known to occur after whatever motif was in current position-1. In so doing, the script will generate many HYP1 HVDs. Finally, this list is triaged to report only those that have >20 motifs, and have known start or end patterns because they seem to be the most conserved arrangements, using a second custom python script (Triage_simulated_HVDs_based_on_known_starts_and_ends.py). One million iterations were performed.

## Author contributions

US performed the experiments and data analysis, HB and BS prepared the biological material, PT, KV, and JTJ were responsible for the sequencing and assembly of the genome of *G. pallida* Newton. SEVDA and US drafted the manuscript. All authors read and approved the final manuscript.

## Availability of data and materials

Raw reads and assembled genomes will be made publicly and freely available upon acceptance of the manuscript: BioProject number XYZ.

## Declarations

### Ethics approval and consent to participate

Not applicable.

### Consent for publication

Not applicable.

---

**Figure S1.**
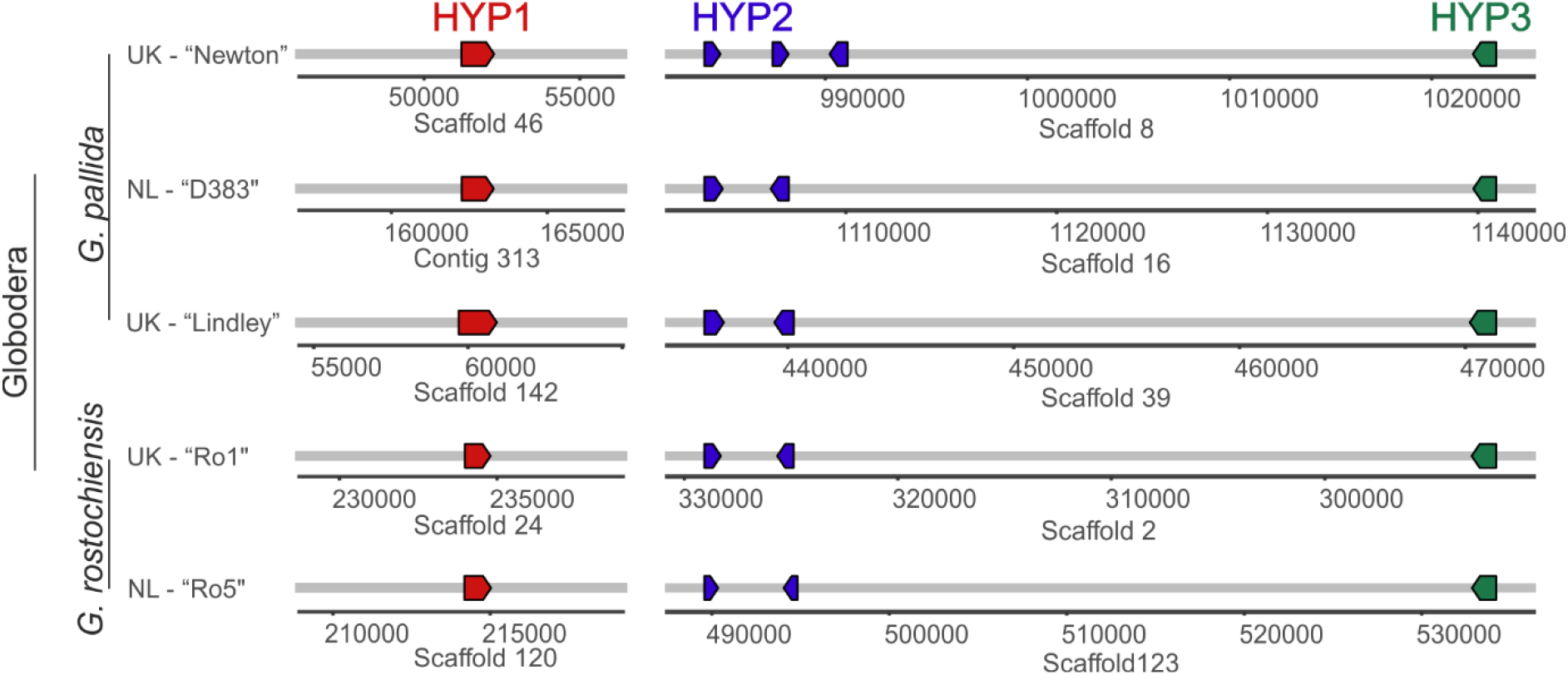
Schematic gene maps indicating positions of HYP1, HYP2 and HYP3 loci across multiple populations/pathotypes of *G. pallida* and *G. rostochiensis*.

**Figure S2.**
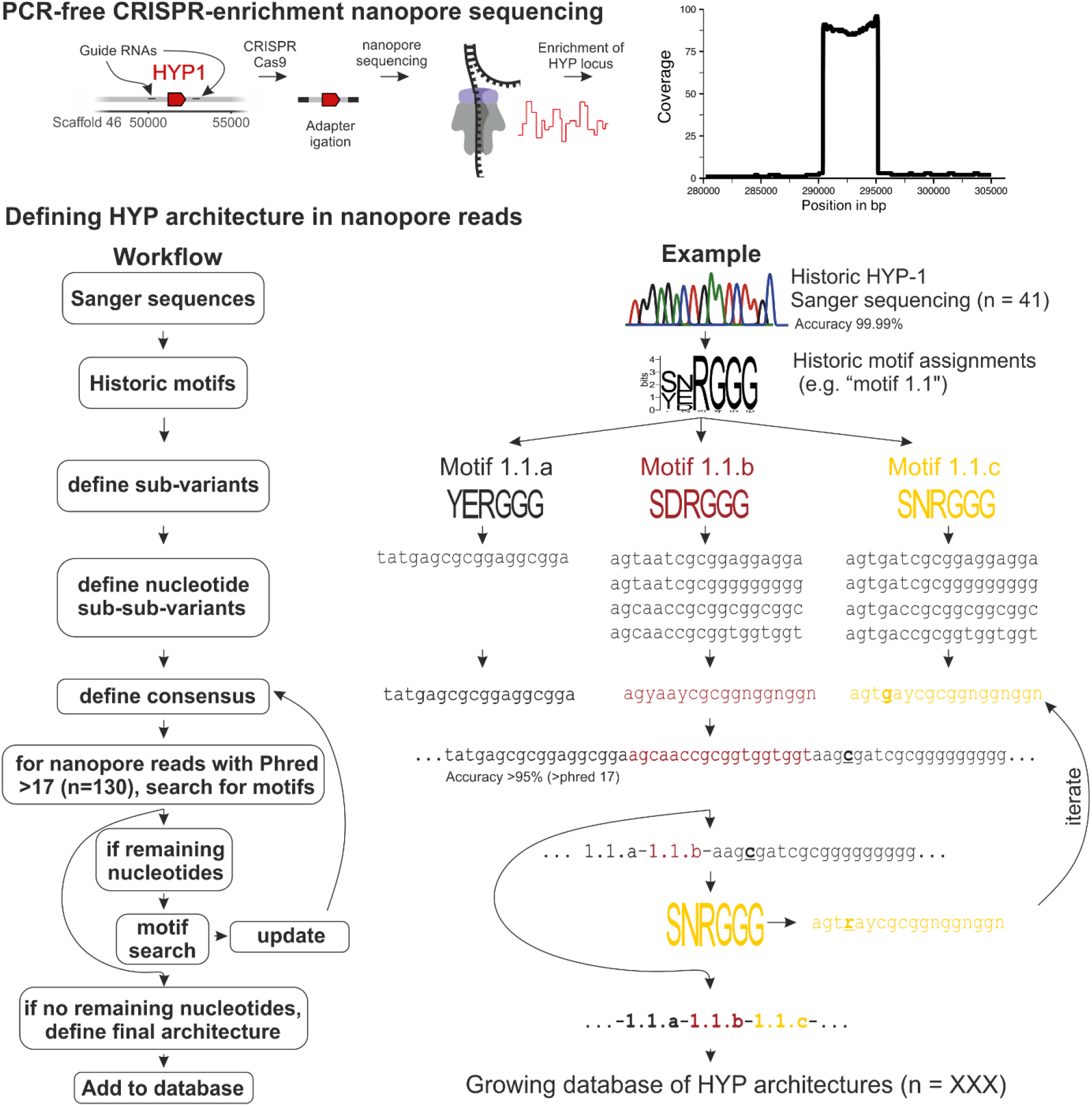
Schematic of workflow for characterization of HYP variants within a population or single nematode. A customised workflow was designed by taking advantage of the ‘modular’ nature of the motifs within the variable region of HYP genes. Previously identified motifs or those identified by MEME were used to build a pattern matching code using the Biostrings package in R. The pattern matching was optimised for error-prone nature of long reads. Several iterations were performed to update code for missed motifs. Finally, known motifs missed due to multiple sequencing errors were manually curated to generate a final list of HYP ‘variants’ consisting of an uninterrupted string of identified motifs.

**Figure S3.**
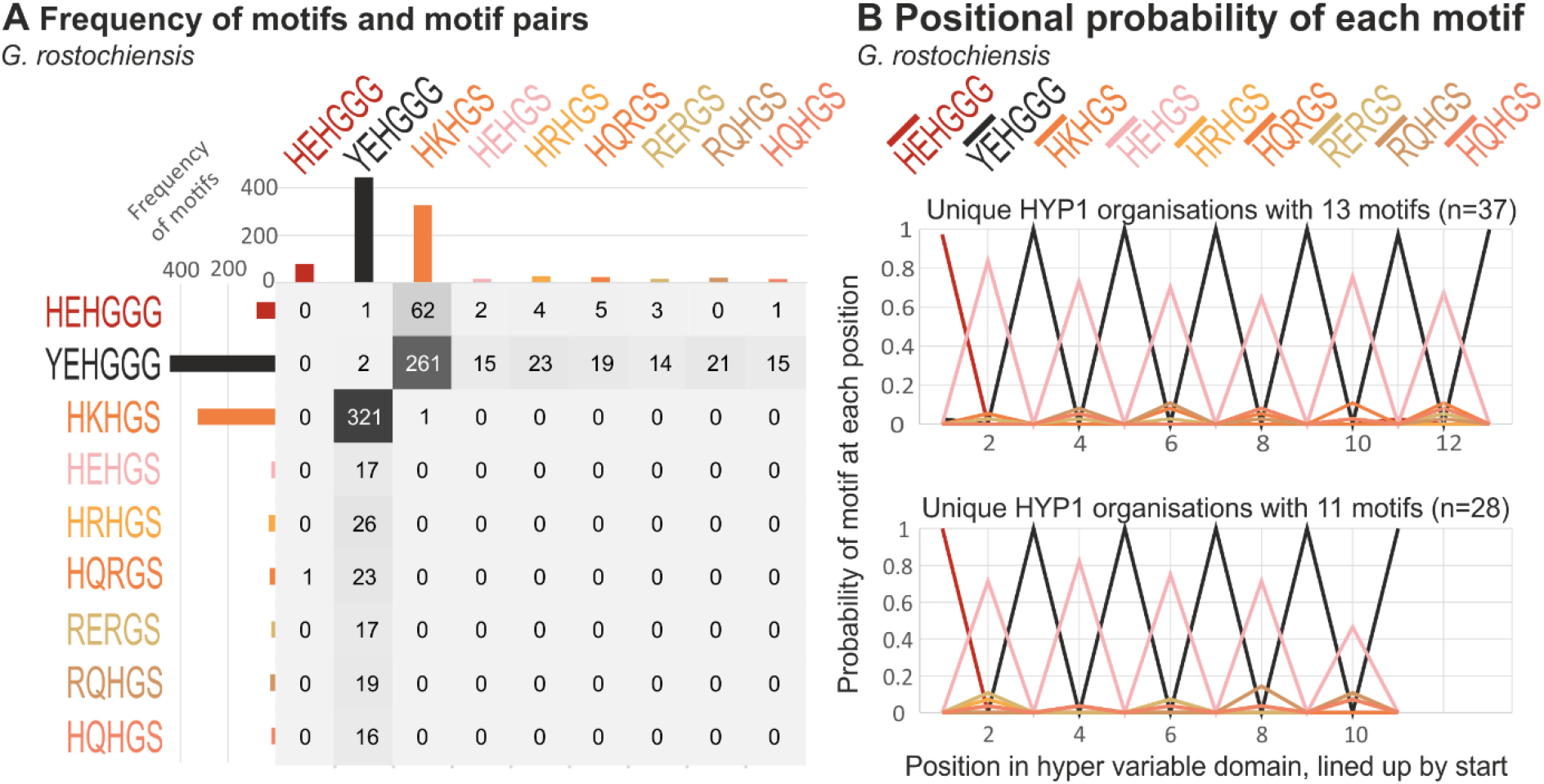
There are different rules underlying *G. rostochiensis* HYP1 variation. A) Frequency of motif pairs within HYP1 HVDs. Each motif is shown with amino acid sequence, y axis of matrix is position n and x axis is position n+1. For examples: i) HEHGGG is almost always first (and so almost never follows another motif), ii) all motifs can appear before YEHGGG, but YEHGGG is the only motif that can appear before RQHGS). B) The positional probability of each motif at each position for HYP1s with HVDs containing 13 (top) and 11 (bottom) motifs show an alternating pattern of YEHGGG followed by an other motif.

**Figure S4.**
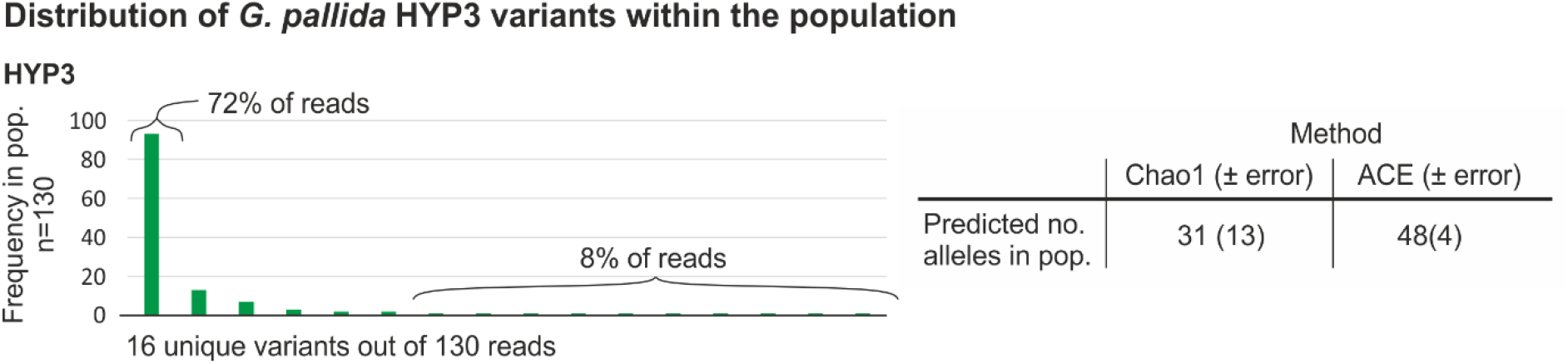
Frequency distribution and population size estimates of *G. pallida* HYP3 alleles. The observed occurrences of each unique HYP variant when sampling a population (n= 125). Inset are two independent species abundance estimates for the total number of alleles in the population, based on the sampling.

**Figure S5.**
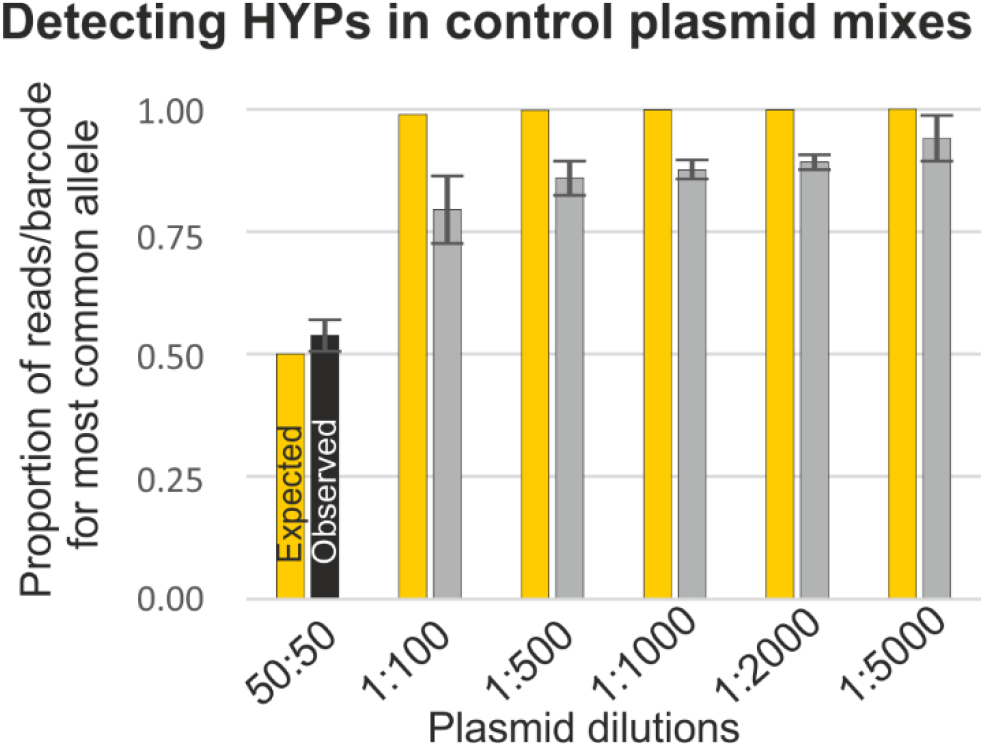
Detecting HYPs in control plasmid mixes. Comparison of observed (black and grey) and expected (yellow) proportions of reads per barcode pair that are contributed by the most common allele, for a range of known plasmid dilutions.

**Figure S6.**
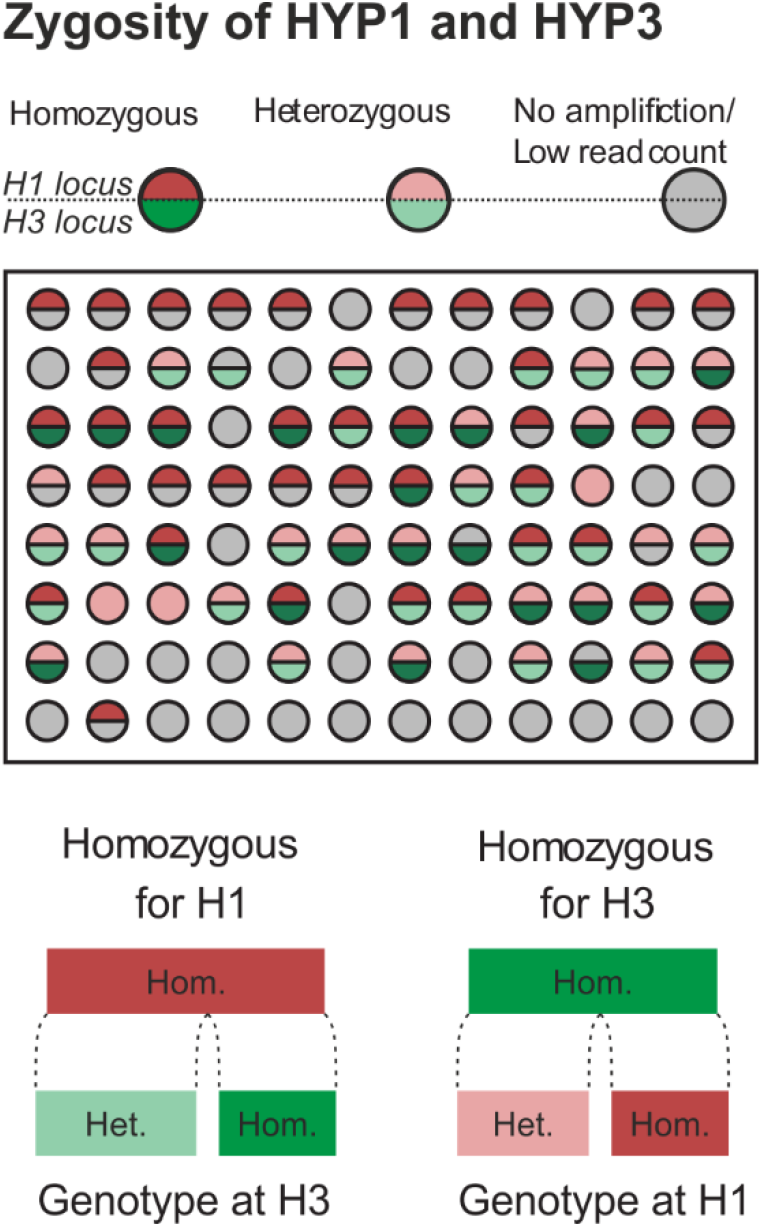
Zygosity of HYP1 and HYP3 loci for 68 J2s. Zygosity of HYP1 locus (reds) is unlinked to zygosity of HY3 locus (greens).

**Table S1:** Genome metrics for Globodera genome assemblies.

| **Metrics** | ***G. pallida* 'Lindley'**<br>**(Cotton et al., 2014)**<br>**shortread assembly** | ***G. pallida* 'Newton' longread**<br>**assembly** | ***G. pallida* 'Lindley'**<br>**longread assembly** | ***G. rostochiensis* 'Ro1'**<br>**(Eves-van den Akker et al., 2016)**<br>**shortread assembly** | ***G. rostochiensis***<br>**'Ro1'**<br>**longread assembly** |
| --- | --- | --- | --- | --- | --- |
| **Size (Mb)** | 124.6 | 119.6 | 109.1 | 95.9 | 102.2 |
| **Scaffolds(n)** | 6,873 | 173 | 672 | 4281 | 267 |
| **Scaffold N50 (bp)** | 121,687 | 2,225,126 | 595,505 | 88,688 | 2,199,277 |
| **Longest scaffold**<br>**(bp)** | 600,076 | 8,303,752 | 4,690,510 | 688,384 | 8,068,213 |
| **GC (%)** | 37 | 37 | 37 | 38 | 39 |
| **Ns (bp)** | 21,024,229 | 1,237,729 | 2,170,753 | 4,399,212 | 1,328,123 |
| **BUSCO (%)** | Complete:73%<br>[Duplicated:13%],<br>Fragmented:12%,<br>Missing:14%, | Complete: 89% [Duplicated:<br>9.9%],Fragmented: 5.9%,<br>Missing: 4.6% | Complete: 86% [Duplicated:<br>12%], Fragmented: 5.2%,<br>Missing: 8.2% | Complete: 88% [Duplicated: 9.5%],<br>Fragmented: 5.9%, Missing: 5.9% | Completed:87%<br>[Duplicated:9.2%],Fragmented<br>:5.9%,Missing:6.6% |
| **Predicted genes**<br>**(n)** | 16,466 | 16,321 | 22,426 | 14,378 | 14309 |

